# Scalable Analysis of Authentic Viral Envelopes on FRONTERA

**DOI:** 10.1101/2020.07.05.188367

**Authors:** Fabio González-Arias, Tyler Reddy, John E. Stone, Jodi A. Hadden-Perilla, Juan R. Perilla

## Abstract

Enveloped viruses infect host cells via fusion of their viral envelope with the plasma membrane. Upon cell entry, viruses gain access to all the macromolecular machinery necessary to replicate, assemble, and bud their progeny from the infected cell. By employing molecular dynamics simulations to characterize the dynamical and chemical-physical properties of viral envelopes, researchers can gain insights into key determinants of viral infection and propagation. Here, the Frontera supercomputer is leveraged for large-scale analysis of authentic viral envelopes, whose lipid compositions are complex and realistic. VMD with support for MPI is employed on the massive parallel computer to overcome previous computational limitations and enable investigation into virus biology at an unprecedented scale. The modeling and analysis techniques applied to authentic viral envelopes at two levels of particle resolution are broadly applicable to the study of other viruses, including the novel coronavirus that causes COVID-19. A framework for carrying out scalable analysis of multi-million particle MD simulation trajectories on Frontera is presented, expanding the the utility of the machine in humanity’s ongoing fight against infectious disease.

## 1 Introduction

Viruses are pathogens that cause infectious diseases in living organisms. Because of their of lack metabolic capabilities, viruses require the molecular machinery of a host cell to replicate [1]. Virus particles, referred to as virions, exhibit chemical and structural diversity across families. The detailed architecture of a virus determines its fitness, mechanism of infection and replication, and host cell tropism.

The most basic virions consist of genomic material protected by a protein shell, called a capsid. The viral genome contains the blueprint for synthesis of the macromolecular components that will form progeny virions. Depending on the virus, the genome may be encoded as RNA or as DNA [2], [3]. More complex virus structures exhibit an exterior membrane, called an envelope, composed of a lipid bilayer. Fusion of the viral envelope with the plasma membrane of the host cell initiates infection. Enveloped viruses typically incorporate surface glycoproteins that interact with host cell receptors to mediate adhesion and facilitate membrane fusion [4]. The composition of the viral envelope, along with the specificity of the adhesion proteins, contributes to the recognition, attachment, and fusion of the pathogen with its host [5], [6]. Other membrane-embedded structures that may be displayed by a virion include viral ion channels, called viroporins, proteins that provide particle scaffolding or assembly support, or proteins that participate in the release of progeny virions [7].

Enveloped viruses, such as influenza A, Ebola, and HIV-1, account for numerous cases of infection and death in humans each year. SARS-CoV-2, the novel coronavirus that causes COVID-19, is also enveloped. The SARS-CoV-2 virion incorporates a class-I fusion glycoprotein called the spike (S), a purported viroporin known as the envelope (E) protein, and a structurally essential integral membrane (M) protein [8]. While each of these components plays a key role in the viral life cycle, it is ultimately the envelope that enables each to carry out its function successfully. As the envelope is derived from the host cell membrane, particularly from the plasma membrane or organelle in which the virion assembles, its lipid composition may be highly complex [9]. Further, the envelope may be asymmetric, or present different lipid compositions across the inner versus outer leaflets of the bilayer; membrane asymmetry is crucial to the function of most organelles and may likewise be important in the assembly or infection processes of viruses [10], [11].

Detailed characterization of viruses is critical to the development of new antiviral therapies [12]. Computational studies now routinely contribute to the foundation of basic science that drives innovation in pharmacy, medicine, and the management of public health. Notably, molecular dynamics (MD) simulations have emerged as a powerful tool to investigate viruses. Following advances in high-performance computing, MD simulations can now be applied to elucidate chemical-physical properties of large, biologically relevant virus structures, revealing insights into their mechanisms of infection and replication that are inaccessible to experiments [13]–[16]. In recent years, MD simulations of intact enveloped virions, including influenza A, dengue, and an immature HIV-1 particle, have been reported. State-of-the art lipidomics profiling of viruses has enabled some of these models to contain realistic lipid composition, leading to the first computational studies of authentic viral envelopes [17]–[20]. Here, next-generation simulation and analysis of authentic viral envelopes is discussed, leveraging the resources of the leadership-class Frontera supercomputer. A framework for deploying large-scale analysis of MD simulation trajectories on Frontera is presented.

## 2 Computational challenges

MD simulations of membranes can be carried out at coarse-grained (CG) or atomistic levels of detail. CG models employ particles that consolidate groups of atoms. Atomistic models employ discrete particles for each constituent atom, including hydrogen. Due to the size and chemical complexity of authentic viral envelopes, MD simulations can include millions – even hundreds of millions – of particles, particularly when the physiological solvent environment of the system is taken into account. Moreover, MD simulations of membranes must be performed on timescales of hundreds of nanoseconds to microseconds in order to characterize dynamical properties, such as lipid lateral diffusion and flip-flop between leaflets of the bilayer [16], [21].

CG models reduce computational expense by eliminating degrees of freedom, enabling the exploration of extended simulation timescales; however, the loss of detail versus atomistic models reduces simulation accuracy. An approach that combines the strengths of both methods involves building and simulating a CG model, which reproduces essential features of the membrane and allows equilibration of lipid species [22], then backmapping the CG representation to an atomistic model for further study [23], [24]. The backmapping process can be precarious and depends on the molecular mechanics force field and energy minimization scheme to resolve structural artifacts. Either way, the incredible number of particle-particle interactions that must be computed to reproduce the behavior of a membrane bilayer over extended simulation timescales, especially for complex, large-scale systems like viral envelopes, presents a significant computational challenge [25].

Furthermore, the amount of data generated by MD simulations of intact viral envelopes is tremendous, both in terms of chemical intricacy and storage footprint. MD simulation trajectories, which may range from tens to hundreds of terabytes in size, fail to provide new information about viruses until they are analyzed in painstaking detail. The magnitude of trajectory files and the sheer numbers of particles that must be tracked during analyses necessitates massively parallel computational solutions. It follows that such large datasets cannot be feasibly transferred to other locations for analysis or visualization, and interaction with the data to yield scientific discoveries must be conducted on the same high-performance computing resource used to run the simulation in the first place.

## 3 Computational solutions on Frontera

Frontera is a leadership-class petascale supercomputer, funded by the National Science Foundation, and housed at the Texas Advanced Computing Center at the University of Texas at Austin. It is ranked as the fifth most powerful supercomputer in the world, and the fastest on any university campus. Frontera provides an invaluable computational resource for furthering basic science research of large biological systems, including enveloped viruses.

Frontera comprises multiple computing subsystems. The primary partition is CPU-only and consists of 8,008 compute nodes powered by Intel Xeon Platinum 8280 (CLX) processors; each node provides 56 cores (28 cores per socket) and 192 GB of RAM. The large memory partition includes 16 additional compute nodes with 112 cores (28 cores per socket) and memory upgraded to 2.1 TB NVDIMM. Another partition is a hybrid CPU/GPU architecture, consisting of 90 compute nodes powered by Intel Xeon E5-2620 v4 processors and NVIDIA Quadro RTX 5000 GPUs; each node provides 16 cores (8 cores per socket), four GPUs, and 128 GB of RAM. All Frontera nodes contain 240 GB solid-state drives and are interconnected with Mellanox HDR InfiniBand. The Longhorn subsystem consists of 96 hybrid CPU/GPU compute nodes powered by IBM POWER9 processors and NVIDIA Tesla V100 GPUs; each node provides 40 cores (20 cores per socket), four GPUs, 256 GB of RAM, and 900 GB of local storage. Eight additional nodes have RAM upgraded to 512 GB to support memory-intensive calculations. Longhorn is interconnected with Mellanox EDR InfiniBand.

Frontera has a multi-tier file system and provides 60 PB of Lustre-based storage shared across nodes. The ability to store massive amounts of data and analyze this data in parallel enables the investigation of biological systems at an unprecedented scale. Researchers gain access, not only to the computational capability to perform MD simulations of millions of particles, but also to the capability to extract more complex and comprehensive information from their simulation trajectories, leading to deeper discoveries relevant to the advancement of human health.

Frontera’s Lustre filesystem is crucial for the performance of large-scale biomolecular analysis. Lustre decomposes storage resources among a large number of so-called Object Store Targets (OSTs) that are themselves composed of moderate sized high-performance RAID-protected disk arrays. In much the same way that a RAID-0 array can stripe a file over multiple disks, Lustre can similarly stripe files over multiple OSTs. This second level of striping over OSTs is particularly advantageous when working with very large files, since I/O operations at different file offsets can be serviced in parallel by multiple independent OSTs, according to the stripe count and stripe size set by the user, or by system defaults. Figure 1A illustrates how an MD simulation trajectory (and potentially even individual frames) can be striped over OSTs. Figure 1B demonstrates the scalability of multi-million particle biomolecular analysis on Frontera when using the Lustre filesystem.

**Fig. 1.**
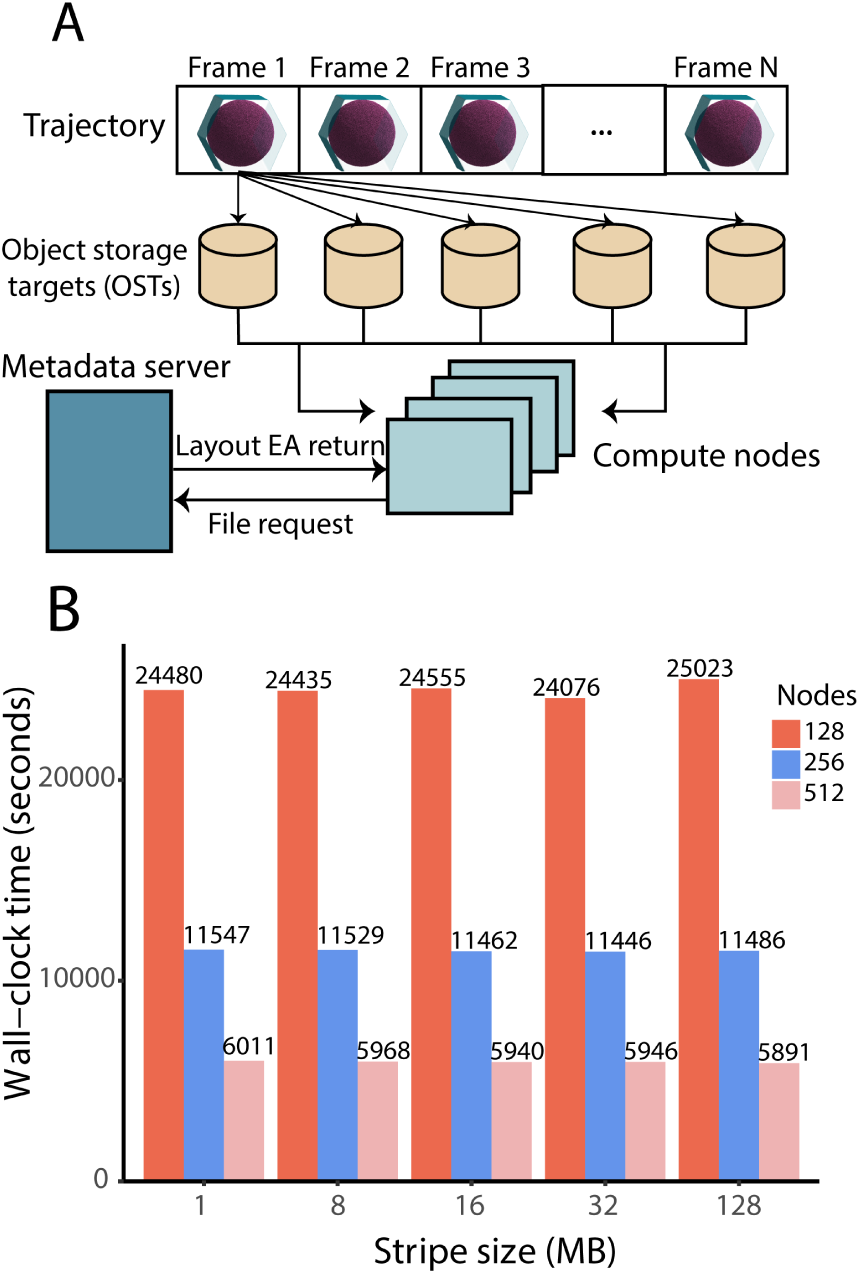
Analysis of large-scale MD simulation trajectories using Lustre filesystem **A** Illustration of trajectory file striping using Lustre filesystem. Each frame on the file is striped across multiple OSTs, where the number of stripes are spawned according to the stripe count and stripe size of the storage file. Upon submission of the job, the compute nodes access a series of frames, from the OSTs, to perform the analysis of the trajectory using parallel I/O. Instructions from the Tcl scripting interface involving reading and writing files are requested through the Metadata Server (MS), fetching the layout extended attributes (EAs) and using this information to perform I/O on the file. **B** Wall-clock time for the transverse diffusion analysis of authentic viral envelopes on Frontera, using a stripe count of 16 with different stripe sizes.

## 4 Parallel Analysis with VMD

Visual Molecular Dynamics (VMD) is a widely-used software for biomolecular visualization and analysis [26]. Commonly utilized as a desktop application to setup MD simulations and interact with trajectory data, VMD exploits multi-core CPUS, CPU vectorization, and GPU-accelerated computing techniques to achieve high performance on key molecular modeling tasks. VMD can also be used in-situ on massive parallel computers to perform large-scale modeling tasks, exploiting computing and I/O resources that are orders of magnitude greater than those available on even the most powerful desktop workstations. MPI implementations of VMD have already been employed on supercomputers to enable novel investigation of large virus structures, such as the capsid of HIV-1 [27].

To facilitate large-scale molecular modeling pipelines, VMD incorporates support for distributed memory message passing with MPI, a built-in parallel work scheduler with dynamic load balancing, and easy to use scripting commands, enabling large-scale parallel execution of molecular visualization and analysis of MD simulation trajectories. VMD’s Tcl and Python scripting interfaces provide a user-friendly mechanism to distribute and schedule user-defined work across MPI ranks, to synchronize workers, and to gather results. This approach allows user-written modeling and analysis scripts to be readily adapted from existing scripts and protocols that have been developed previously on local computing resources, allowing robust tools to be deployed on a large-scale, often with few changes.

VMD exploits node-level hardware-optimized CPU and GPU kernels, all of which are also used when running on distributed memory parallel computers. Computationally demanding VMD commands are executed by the fastest available hardware-optimized code path, with a general purpose C++ implementation as the standard fall-back. When run in parallel, the I/O operations of each VMD MPI rank are independent of each other, allowing I/O intensive trajectory analysis tasks to naturally exploit parallel filesystems.

VMD supports I/O-efficient MD simulation trajectory formats that have been specifically designed for so-called burst buffers and flash-based high-performance computing storage tiers, with emerging GPU-Direct Storage interfaces achieving I/O rates of up to 70 GB/sec on a single GPU-dense compute node, and conventional parallel I/O rates approaching 1 TB/sec. VMD startup scripts can be customized to run one or multiple MPI ranks per node, while avoiding GPU sharing conflicts or other undesirable outcomes, allowing compute, memory, and I/O resources to be apportioned among MPI ranks, and thereby best-utilized for the task at hand.

To facilitate large-scale simulation and analysis of biological systems containing millions of particles, VMD has been compiled on Frontera with MPI enabled. Frontera’s capacity for massively parallel investigation of intact, authentic viral envelopes is demonstrated by application to envelope models at two levels of particle resolution.

## 5 Modeling authentic viral envelopes

### 5.1 Coarse-grained model of HIV-1 envelope

Lipidomics profiling by mass spectrometry has established the lipid composition of the HIV-1 viral envelope [28], [29]. Based on this experimental data, a CG model of an authentic HIV-1 envelope was constructed (Fig. 2A). The model exhibits a highly complex chemical makeup, with 24 different lipid species and asymmetry across the inner and outer leaflets of the membrane bilayer. The envelope is 120 nm in diameter and, with solvent, comprises 29 million CG beads, represented as MARTINI [30] particles. The CG model was equilibrated using GROMACS 2018.1 [31], and its dynamics were investigated over a production simulation time of over five microseconds. The MD simulation system, complete with solvent environment, is shown in Figure 2B.

**Fig. 2.**
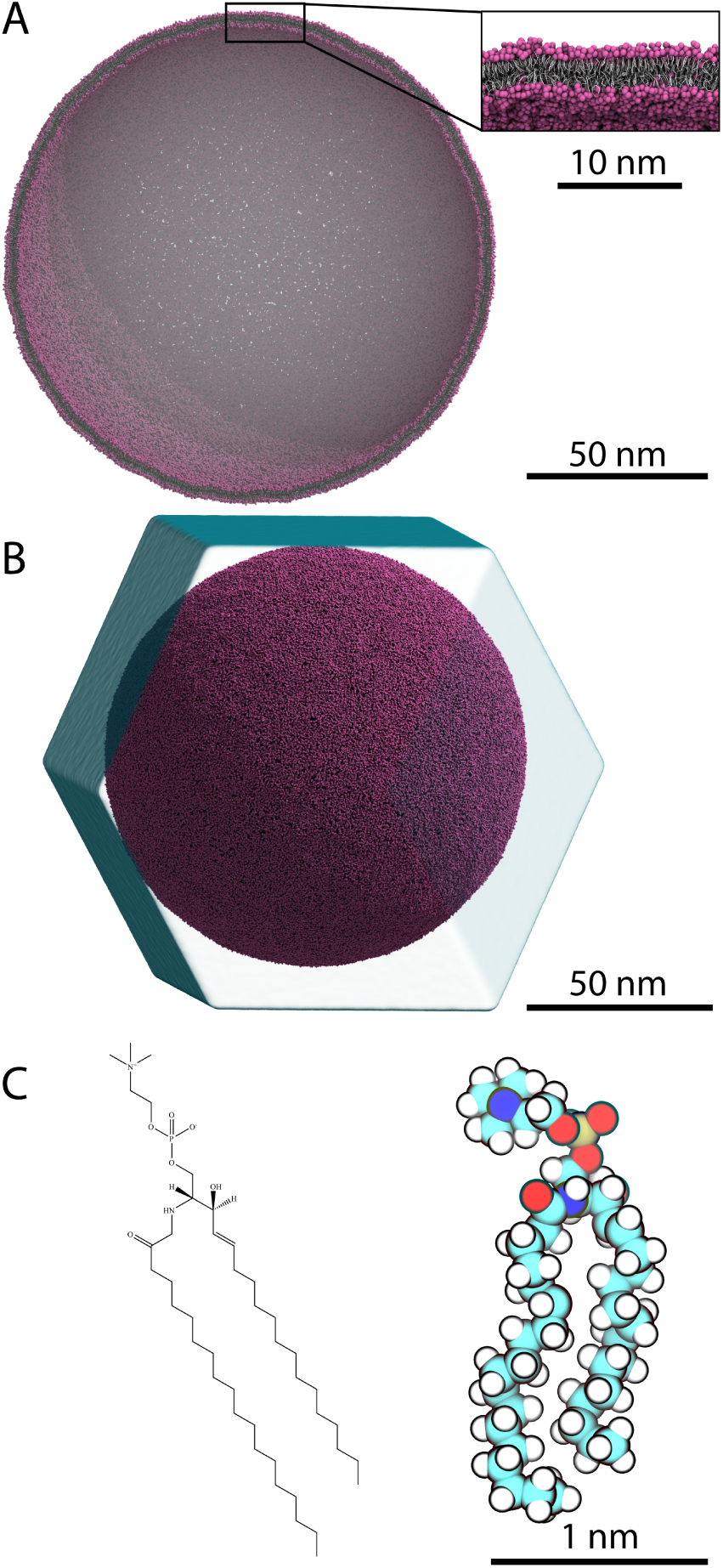
Authentic model of HIV-1 viral envelope **A** Cross-section of CG envelope model. Inset shows details of the inner (endoplasmic) and outer (exoplasmic) leaflets of the membrane bilayer. Envelope is 120 nm in diameter and comprises 29 million MARTINI particles. **B** CG envelope model suspended in physiological solvent environment, used for long-timescale MD simulations. **C** Chemical structure and atomistic representation of DPSM, one of 24 lipid species included in the HIV-1 envelope model. Each CG lipid in the envelope was backmapped to an atomistic version exhibiting full chemical detail.

### 5.2 Atomistic model of HIV-1 envelope

All-atom MD simulations are the most accurate classical mechanical approach that can be applied to study largescale biological systems [13]. Following simulation of the CG HIV-1 envelope model, which facilitated equilibration of lipid distributions over an extended simulation timescale, an atomistic model was constructed based on backmapping. The equilibrated CG model was mapped to an atomistic representation using the Backward tool [24], which enables transformation of CG systems built with MARTINI [30]. The approach utilizes a series of mapping files that contain structural information and geometric restrictions relevant to the modeling of each lipid species. Local geometries of atomistic lipids are reconstructed, taking into account stoichiometry and stereochemistry, to project the CG configuration into an atomistic configuration. An example of an atomistic representation of a lipid, along with its chemical structure, is given in Figure 2C. The final atomistic model of an authentic HIV-1 viral envelope comprises over 280 million atoms including solvent and ions. To complete the backmapping process, the system must be relaxed to a local energy minimum to resolve structural artifacts introduced by the CG to all-atom transformation.

## 6 Viral envelope analysis on Frontera

### 6.1 Measuring transverse lipid diffusion

Transverse diffusion of lipids occurs when they flip-flop between leaflets of the membrane bilayer. Cellular enzymes can catalyze this movement of lipids, or it can occur spontaneously over longer timescales. Computational approaches to characterize transverse diffusion in heterogeneous lipid mixtures have been described previously and indicate that flip-flop occurs at relatively slow rates [32], [33]. Current strategies to measure flip-flip events are based on tracking the translocation and reorientation of lipid headgroups (i.e., colored beads in Fig. 2A).

A feature released in VMD 1.9.4 by the authors (*measure volinterior*) [34] facilitates tracking of headgroup dynamics by providing rapid classification of their locations in the inner versus outer leaflets of the bilayer. The method equips VMD with a mechanism to identify the inside versus outside of a biomolecular container. If the container surface is specified as the region of the bilayer composed of lipid tail groups (i.e., silver in Fig. 2A), VMD can detect the presence of lipid headgroups in the endoplasmic versus exoplasmic leaflets. By tracking the number of headgroups that flip-flop from one leaflet to the other during intervals of simulation time, rates of transverse diffusion can be readily calculated [34].

Frontera was leveraged to measure transverse lipid diffusion in the CG model of the authentic HIV-1 viral envelope using *measure volinterior* in VMD compiled with MPI support. The entire 5.2 *μ*s of MD simulation, comprising 5,200 frames with a storage footprint of 170 GB, was used for the calculation. To obtain high I/O performance, the trajectory files were set to a 1 MB stripe width, striping over a total of 16 Lustre OSTs. The analysis was run on Frontera’s primary compute partition, utilizing 56 Intel CLX cores per node.

Flip-flop events for the lipid DPSM were found to occur at a rate of 2.62 ×10^−4^ ± 1.49 × 10^−5^ molecules per nanosecond from the exoplasmic to endoplasmic leaflet, and 2.33 × 10^−5^ ± 1.41 × 10^−6^ molecules per nanosecond from the endoplasmic to exoplasmic leaflet (Fig. 3A). Because inward versus outward diffusion of this lipid species is not occurring at the same rate, the calculation reveals the process of lipid distributions being equilibrated across the asymmetric bilayer over the course of the simulation. The Tcl framework for performing the calculation with VMD, the VMD startup script, and the SLURM job submission script used to run the calculation on Frontera, are given in the Supplementary Information.

**Fig. 3.**
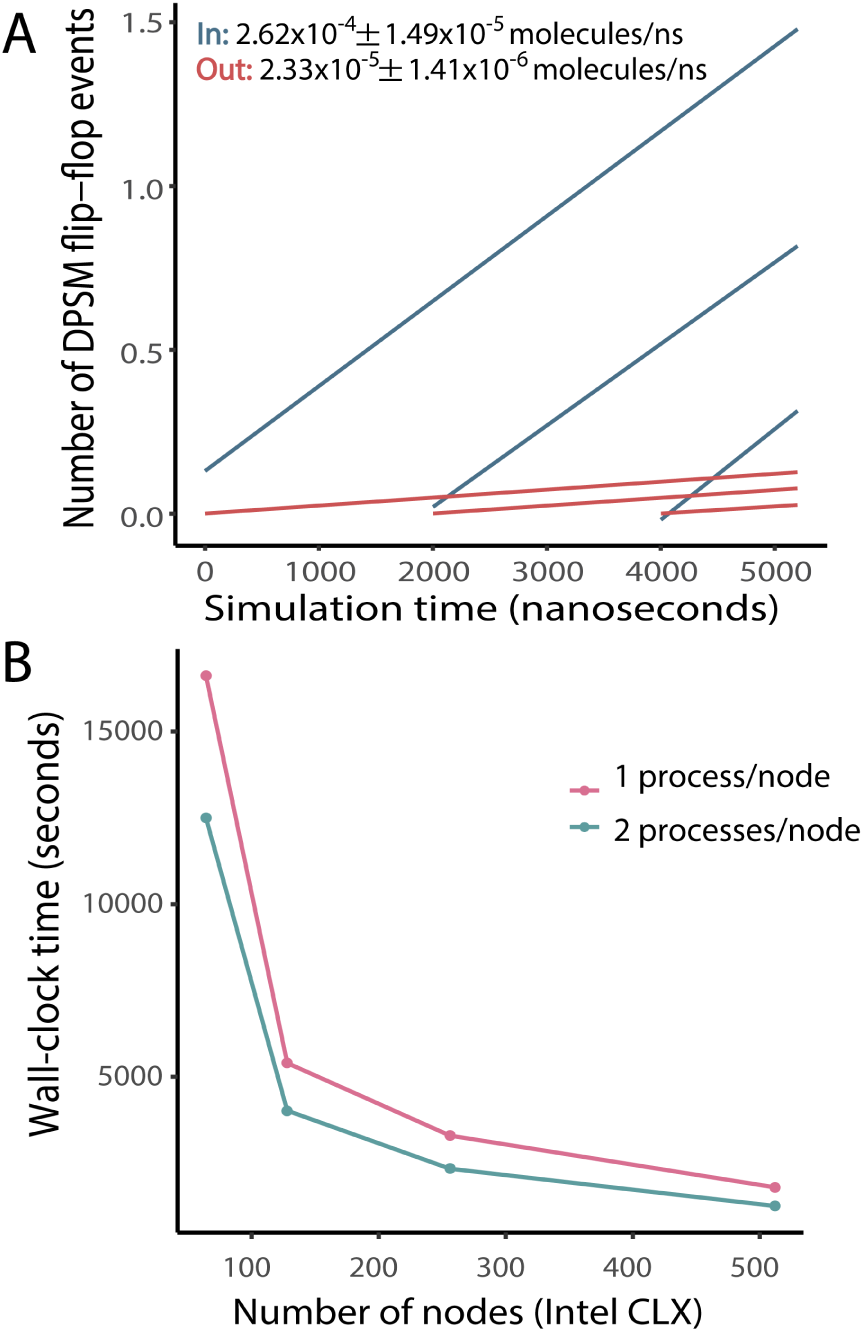
Parallel analysis of authentic viral envelopes. **A** Transverse diffusion analysis of the lipid DPSM in the CG model of the viral envelope. **B** Wall-clock time for calculation of transverse diffusion on Frontera.

Figure 3B shows the wall-clock time for the transverse diffusion calculation with one VMD MPI rank per node and two VMD MPI ranks per node. By running multiple VMD MPI ranks per node, I/O operations are more effectively overlapped with computations, thereby achieving better overall I/O and compute throughput both per-node, and in the aggregate, leading to an overall performance gain. It is worth noting, however, that finite per-node memory, GPU, and interconnect resources ultimately restrict the benefit of using multiple MPI ranks per-node to achieve better compute-I/O overlap. Ongoing VMD developments aim to improve compute-I/O overlap both for CPUs and for GPU-accelerated molecular modeling tasks through increased internal multithreading of trajectory I/O to allow greater asynchrony and decoupling of I/O from internal VMD computational kernels.

### 6.2 Iterative relaxation of an atomistic envelope

The atomistic model of an authentic HIV-1 viral envelope presented here was derived via backmapping from a CG model. A major limitation of the backmapping approach involves structural artifacts introduced by the CG to all-atom transformation. Figure 4A shows an example of such structural artifacts in the lipid DPSM, including distorted bond lengths and angles. The current strategy for remedying these artifacts is to subject the backmapped model to molecular mechanics energy minimization, relaxing the geometry of each molecule, and the system as a whole, to a local energy minimum. Figure 4B shows an example of a post-minimization structure in which the distortions have been successfully resolved.

**Fig. 4.**
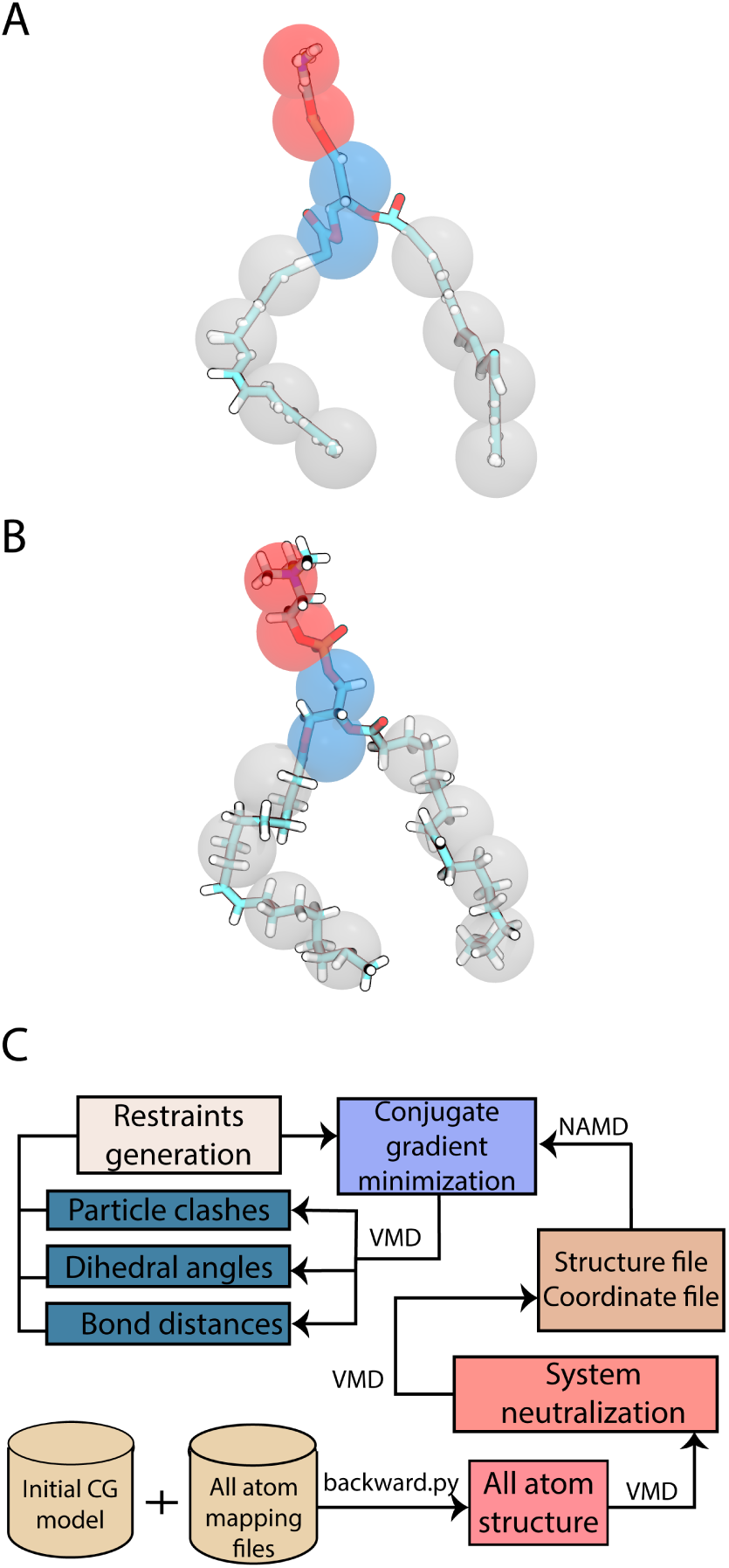
Resolving distortions in backmapped lipids. **A** Atomistic representation of DPSM with structural artifacts post-backmapping. CG representation of the lipid is superimposed to the all-atom structure. **B** Atomistic representation of DPSM post-minimization with structural artifacts resolved. CG representation of the lipid is superimposed to the relaxed all-atom structure. **C** Iterative relaxation process for large-scale authentic atomistic viral envelopes. Following backmapping from CG representations to all-atom, the system is subjected to charge neutralization, followed by generation of topology and coordinate files. Conjugate gradient minimization using NAMD, in combination with parallel analysis using VMD MPI, is employed iteratively to eliminate structural artifacts of the membrane and drive the system to a local minimum.

Energy minimization is a common procedure applied to biomolecular systems to eliminate close contacts and non-ideal geometries prior to the start of MD simulations. The energy of atomistic systems is described as an empirical potential energy function *V(r*_*i*_), where the terms depend on the positions *r* of *i* atoms [35]. Here, energy minimization of the system is performed on Frontera using NAMD 2.14, an MD engine based on C++ and charmm++ that is amenable to large systems by virtue of being highly scalable on massive parallel computers [36], [37]. Energy minimization is implemented in NAMD using the conjugate gradient algorithm [38] for maximum performance. The atomistic model of an authentic HIV-1 viral envelope was subjected to energy minimization based on the CHARMM36 force field, with all the latest corrections in parameters for lipids and heterogenous protein-lipid systems [39].

Figure 4C diagrams the complete procedure for constructing an atomistic envelope model from an initial CG model. Following backmapping of the CG representation to an all-atom representation, explicit ions must be added to achieve electrostatic neutralization of the system, given the numerous charged lipid headgroups present in the envelope. Coordinate and topology files are generated using the *js* plugin in VMD, utilizing a binary file format that overcomes the fixed-column limitation of *pdb* files. The minimization of multi-million atom systems can occasionally cause memory overflow or significant loss of performance in NAMD, due to the I/O, even on large-scale computational resources. In remedy, the memory-optimized version of NAMD implements parallel I/O and a compressed topology file, reducing the memory requirements for large biomolecular systems.

Following initial conjugate gradient minimization with NAMD, the model is assessed for major contributors to energetic instability, including structural clashes, close contacts, bond lengths or angles significantly exceeding equilibrium values, and unfavorable dihedral configurations. Such assessments can be rapidly accomplished using VMD compiled with MPI support on Frontera, using existing VMD commands like *measure contacts, measure bond, measure angle*, and *measure dihed*. In a second iteration of minimization, regions of the system that are relatively stable are restrained in order to focus further minimization efforts on regions of that remain the most energetically unfavorable. Useful NAMD commands for minimization of unstable systems include *minTinyStep* and *minBabyStep*, which control the magnitude of steps for the line minimizer algorithm for initial and further steps of minimization, respectively. This process is repeated until all geometric distortions are resolved, and the atomistic model has been driven to a local energy minimum, representative of the stable biological system at equilibrium.

## 7 Conclusions

Increased computational capability presents the opportunity for researchers to push the envelope and extend the bounds of scientific knowledge. Building, simulating, and analyzing models of intact, authentic viral envelopes, particularly at atomistic detail, is a relatively new research endeavor under active development. The leadership-class Frontera supercomputer provides a powerful resource to support the continued growth of this field. The combination of Frontera’s state-of-the-art, massively parallel architecture and high-performance software applications, such as VMD, that can exploit it will enabling researchers to make discoveries relevant to the treatment and prevention of disease at an unprecedented scale. All of the techniques discussed here are applicable to modeling and investigation of other enveloped viruses, including SARS-CoV-2, which likewise comprises a complex lipidome. Owing to resources like Frontera, the computational biological sciences will play an increasingly important role in driving future innovations that improve human health and longevity.

## Acknowledgments

This work is supported by National Science Foundation award MCB-2027096, funded in part by the Delaware Established Program to Stimulate Competitive Research (EP-SCoR). This research is part of the Frontera computing project at the Texas Advanced Computing Center. Frontera is made possible by National Science Foundation award OAC-1818253. VMD development has been supported by National Institutes of Health grant P41-GM104601. This research used resources provided by the Los Alamos National Laboratory Institutional Computing Program, which is supported by the U.S. Department of Energy National Nuclear Security Administration under Contract No. 89233218CNA000001.

## Notes

### Competing Interest Statement

The authors have declared no competing interest.

