## Supplementary information for "Scalable Analysis of Authentic Viral Envelopes on FRONTERA"

#### Contents

**1 VMD MPI startup script** 2

**2 Frontera job submission script** 3

**3 Lipid transverse diffusion script** 4

### 1 VMD MPI STARTUP SCRIPT

**Code Snippet 1: Startup VMD MPI script on Frontera to force CPU count to 56 cores per node**

```
#!/bin/csh
set defaultvmddir="yourdirectory"
set vmdbasename=vmd
...
# Force the number of CPU count
# to the total number of cores per node
# The primary system on Frontera contains 56 cores/node
  @ cpucount = 56
  setenv WKFORCECPUCOUNT  $cpucount
  setenv VMDFORCECPUCOUNT $cpucount
  setenv VMDFORCECPUCOUNT $cpucount
  setenv RTFORCECPUCOUNT  $cpucount

# Set the total number of nodes for the MPI job
  @ nnodes = 512 # The Limit of node request for a single job on Frontera
#set the number of processes per node
  @ procs = 1
  @ totalprocs = $nnodes + $procs
  echo "Launching VMD MPI..."

# TACC-specific MPI launcher ibrun executes VMD MPI
# Using $nnodes nodes and distributing one rank per node
# If more than one process per node is desired, increment
# the variable $procs
  ibrun -n $totalprocs -N $nnodes -bl numanode -bp scatter "$MASTERVMDDIR/$execname" $*
```

#### 2 FRONTERA JOB SUBMISSION SCRIPT

**Code Snippet 2: Submission script to execute VMD MPI on Frontera using Slurm**

```
#!/bin/bash
if ( $# < 3 ) then
  echo "The following submission script requeres 2 arguments:"
  echo "  VMD input file"
  echo "  VMD log file"
   exit -1
endif

#Arguments to execute the job
INPUTFILE=${1}
LOGFILE=${2}

# The following options can be modified according to the running
# time, allocation to use, number of nodes and processes per node
TIME=02:00:00
QUEUE=normal
HOSTS=512
PROCSPERNODE=1
# Specify the running directory and directory where VMD MPI is
# installes
RUNDIR=`pwd`
BINDIR=yourdirectory

sbatch  <<ENDINPUT
#!/bin/bash
#SBATCH --job-name=VMD
#SBATCH --nodes=$HOSTS
#SBATCH --ntasks=$HOSTS
#SBATCH --time=$TIME
#SBATCH --ntasks-per-node=$PROCSPERNODE
#SBATCH --partition=$QUEUE
#SBATCH -A youraccount
#SBATCH -o slurm%j.out
#SBATCH -e slurm%j.err

cd $RUNDIR
# Execute the job using ibrun
ibrun $BINDIR/vmd -e ${INPUTFILE} > ${LOGFILE}
ENDINPUT
```

#### 3 LIPID TRANSVERSE DIFFUSION SCRIPT

**Code Snippet 3: Tcl scripting example to compute bilayer transverse diffusion of lipids using *measure volinterior***

```
# Preparation procedure for parallel analysis:
# Set bounds on nodes, based on framecount and nodecount,
# assign a block of frames to each MPIrank,
# return the start and end frames for each rank.
proc blockdecompose { framecount } {
        set noderank [parallel noderank]
        set nodecount [parallel nodecount]
        set start [expr round ($noderank * $framecount / $nodecount)]
        set end [expr round (( $noderank +1) * $framecount / $nodecount) - 1]
        return [list $start $end]
}

proc gathernodenames { mycount trajectoryfile } {
        set noderank [ parallel noderank ]
        # Only print messages on node 0
        if { $noderank == 0} {
                set fileb [ open "${trajectoryfile}_${noderank}_volIn.dat" w]
        }

# Do a parallel gather resulting in a list of all of the node names
        set datalist [parallel allgather $mycount]

# Save line by line
        if {$noderank == 0} {
                foreach dataline $datalist {
                        foreach line $dataline {
                                puts $fileh $line
                        }
                }
        close $fileb
        }
}

# Analysis procedure:
# call the preparation ( blockdecompose ) procedure,
# wait for each node to gather its information,
# load the trajectory from striped directory,
# each node loops over its frames and runs the analysis
proc mpianalyze { framecount trajectoryfile molid } {
        set noderank [parallel noderank]
        set nodecount [parallel nodecount]
        set block [blockdecompose $framecount]
        set start [lindex $block 0]
        set end [lindex $block 1]
        set len [expr $end - $start + 1]

        parallel barrier ; # wait for all nodes to reach this point
        mol addfile $trajectoryfile first $start last $end waitfor all
```

```
        # Wait for all nodes to reach this point
        parallel barrier
        set mydata [list ]

        # Wait for all nodes to reach this point
        parallel barrier

 # Parameters  for  measure  volinterior
 # !! Must be parameterized on a per-system basis, see Bryer et al, JCIM, 2019.
        set  RadiusScale 1.6
        set  Isovalue 2.0
        set  GridSpacing 2.33
        set  Nrays 64

 # Initialize lists
        set lipidtypes [list CHOL DPCE DPSM PGPE PIPS POPE PQPS PRPS DLPC DPGS PAPE PGPS PNSM POPS
                PAPS PIPE POPC PQPE PRPE PUPE]
        set filehList [list ]

foreach lipidtype $lipidtypes {
                file mkdir yourdirectory/$lipidtype
                set idxfile "yourdirectory/${lipidtype}/flip_idx_${lipidtype}_${noderank}.dat"
                set fileh [open $idxfile w]
                lappend filehList $fileh
        }

# Atom selection
        set tails [atomselect $molid "name C4B C5B C6B C4A C5A D4B"]
        $tails set beta 1.00
        set sel1 [atomselect $molid "beta == 1.00" frame 0]

# Setup  macros
        atomselect macro heads "not (not ((name NC3 NH3 CNO PO4 AM1 AM2 GL1 GL2 ROH) or (resname DP

# Classify initial space
         set  Nvoxels [measure volinterior $sel1 -res [expr $RadiusScale / 0.1] -isovalue $Isovalue
                -spacing $GridSpacing -nrays $Nrays -mol $molid -verbose -probmap -count_pmap -disc
# Clean up and release memory
        $sel1 delete
        set stepcycle 10
# Loop  over  trajectory
        for {set frame 0} {$frame <= [expr $len -1]} {incr frame} {
                set ts [expr $frame + $start]
    #Classify space and ovwerwrite
                if { $frame  % 5 == 0 } {
                        set sel [atomselect $molid "beta == 1.00" frame $frame]
                        set  Nvoxels [measure volinterior $sel -res [expr $RadiusScale / 0.1] -isov
                             -spacing $GridSpacing -nrays $Nrays -mol $molid -overwrite 0 -verbose
                        $sel delete
                }
```

```
    # Select head groups
                foreach fileh $filehList lipidtype $lipidtypes {
                        set  LipidsIn [atomselect $molid "same residue as (resname ${lipidtype} and
                        set  LipidsOut [atomselect $molid "same residue as (resname ${lipidtype} an

    # Get  indices  of  lipids  of  inner/outer  leaflet
                        set  LipidsInner [lsort -unique [$LipidsIn get residue]]
                        set  LipidsOuter [lsort -unique [$LipidsOut get residue]]
                        puts $fileh [list $ts $LipidsInner $LipidsOuter]

    # Clean  up and  release  memory
                        $LipidsIn  delete
                        $LipidsOut  delete
                }
# End of loop over frames
        }
# Close output files
        foreach fileh $filehList {
        close $fileh
        }
# Wait for all nodes to reach this point
        parallel barrier
        set mydata $frame
        return $mydata
}

proc analyzetest { mytraj framecount } {
# Wait for all nodes to reach this point
        parallel barrier
        set molid [mol new "mystructurefile.psf"]
# Wait for all nodes to reach this point
        parallel barrier
        set mycount [mpianalyze $framecount $mytraj $molid]
# Wait for all nodes to reach this point
        parallel barrier
        gathernodenames $mycount $mytraj
}
# Execution commands
set mytraj mytrajectoryfile.dcd
set framecount [molinfo $molid get numframes]
analyzetest $mytraj $framecount
quit
```
